# Evaluating artesunate-amodiaquine deployment, efficacy and safety: an *in silico* pharmacological model

**DOI:** 10.1101/567008

**Authors:** Ki Bae Hong, Ian Hastings, Katherine Kay, Eva Maria Hodel

## Abstract

**Background:** The World Health Organization currently recommends artesunate-amodiaquine (AS-AQ) as a first-line treatment for uncomplicated falciparum malaria. The clinical efficacy of AS-AQ is very high but its effectiveness in the field varies considerably. This study aimed at comparing the efficacy, effectiveness and safety of AS-AQ fixed dose combination (FDC) and non-fixed formulation (non-FDC) in controlled and real-life settings using a pharmacological model of antimalarial treatment.

**Methods:** The effectiveness and safety of different drug formulations in different treatment scenarios were investigated using a pharmacological model of AS-AQ treatment. The model simulated multiple treatment scenarios to assess the effects of age-or weight-based dosing bands in three geographically distinct patient populations, and poor patient adherence.

**Results:** The model output was consistent with clinical trials in terms of cure rates, recrudescence rates and the pattern of AQ overdosing with age- and weight-based dosing regimens. AS-AQ treatment has good efficacy and effectiveness in fully adherent patients but monotherapy of AS or AQ lead to treatment failure. The weight-based dosing regimen with FDC was the best option for patients in terms of drug safety and had similar efficacies to the other regimens. Asians were more likely to be overdosed with AQ when using age-based dosing regimens.

**Conclusions:** Weight-based dosing is optimal but not always feasible, so age-based dosing regimens are often used as an alternative. The model outputs highlight the importance of optimising these age-based dosing regimens for specific regions, and identify an increased risk of overdosing in young children.

## Introduction

The World Health Organization (WHO) estimated there were approximately 198 million cases of malaria and 584,000 deaths in 2013 (1). Due to the emergence of chloroquine (CQ), mefloquine (MQ) and sulphadoxine-pyrimethamine (SP) resistance in malaria endemic regions, treatment policy from WHO has been recommending artemisinin-based combination therapy (ACT) as a first-line treatment for uncomplicated *Plasmodium falciparum* malaria for almost a decade (2). Artemisinin derivatives are very potent but rapidly eliminated from the body and should therefore not be administered as monotherapies but in combination with a slower acting partner drugs able to sustain sufficient parasiticidal concentrations to clear all remaining parasites (2). Amodiaquine (AQ) is often used as a seasonal malaria chemoprevention together with SP in children less than five years old who live in high seasonal malaria transmission countries (3). Artesunate-amodiaquine (AS-AQ) is one of the ACTs recommended by the WHO and was adopted as the first-line treatment in West Africa approximately ten years ago (4–7). Recently, AS-AQ was also recommended as the first-line treatment for mass drug administration (MDA) in Ebola-affected countries (8, 9).

Studies of the WHO’s three-day AS-AQ regimen found it to be highly efficacious and safe in the treatment of uncomplicated falciparum malaria (10–16). However, while the clinical efficacy of AS-AQ is high, its real-life effectiveness varies substantially; studies in Africa have shown AS- AQ effectiveness to be between 63-85% (17–22). Many factors affect drug effectiveness in the field, including poor access to treatment, provider compliance to treatment guidelines, or adherence of patients and caregivers to prescriptions (23). AS-AQ can be prescribed as either fixed-dose combination (FDC) formulations or non-FDC (i.e. loose tablets or co-blistered tablets). FDCs include both AS and AQ in a single table, each containing either 25/67.5, 50/135 or 100/270 mg AS/AQ depending on tablet strength (24). While non-FDC co-blister packs contain 50 mg of AS and 153 mg of AQ (25). Non-FDC formulations tend to be less user-friendly and allow patients to take only one of the two ACT components. FDC formulations reduce this problem and are recommended to increase adherence, thus delaying the development of resistance to both drugs (26–29).

It is recommended that the dose of AS-AQ is calculated according to a patient’s body weight (30) but this can be logistically challenging in developing countries. Some health facilities for example may not have functional scales (10, 31). In practice, the amount of drug given to patients is commonly based on their age, which can lead to inaccurate dosing.

Clinical trials are mainly designed to test safety and efficacy of interventions, so the results of trials provide guidance for diagnosis, treatment and prevention. However, conducting clinical trials is expensive, time-consuming and restricted by ethical constraints, particularly when their aim is to directly investigate the impact of poor patient adherence. Pharmacological models of drug treatment can overcome these problems and allow researchers to investigate drug effectiveness and drug regimen safety. Historically, pharmacological models of antimalarial drug treatment focused on monotherapies (32–37) but since the WHO recommendation of ACTs, the model methodologies have been extended to include combination therapies (38–44). This study used a pharmacological model for AS-AQ treatment to compare the efficacy, effectiveness and safety of different dosing regimens and FDC versus non-FDC in controlled and real-life settings. Moreover, this study evaluated the change of drug efficacy and effectiveness when patients adhere poorly to the recommended regimen.

## Methods

### Pharmacological model

This study adapted the pharmacological model described in Kay and Hastings (41) for the simulation of AS-AQ treatment outcome. The model, implemented in the statistical software R (version 3.1.0), tracks the number of parasites *P* over time *t* after treatment using the following differential equation (Equation 10 in (41))

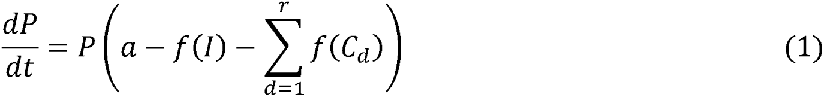

where *P* is the number of parasites, *a* is the parasite growth rate, *f(I)* is the host’s background immunity to the infection (which is assumed to kill parasites or slow their growth), *r* is the number of drugs and/or their active metabolites in the regimen and *f(C_d_)* describes the drug effect (i.e. its kill rate) for each drug *d* depending on its concentration *C_d_* (mg/L). Integrating Equation 1 gives the number of parasites *P* at time *t* (Equation 16 in (41))

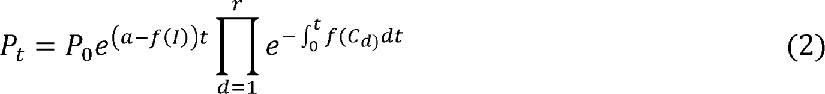

where *P_t_* is the number of parasites at time *t*, *P*_0_ is the number of parasites at the start of treatment. In this study, the hosts’ background immunity was ignored and *f(I)* was set to zero so that all patients were malaria naive and had no acquired immunity.

The drug-dependent killing function for each drug *d* is described by the standard Michaelis-Menton equation, i.e.

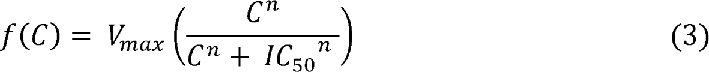

where *V_max_* is the maximal drug-killing rate, *n* is the slope of the dose response curve, and *IC_50_* is the concentration at which 50% of the maximal killing rate occurs.

For AS and its active metabolite dihydroartemisinin (DHA), the model tracked drug concentration over time using a standard one-compartment disposition model allowing for the absorption of the parent drug (i.e. AS) across the gut wall at rate *k_a_* and conversion to its active metabolite DHA at rate *k_m_* (**Figure 1**). Equations for tracking drug concentration over time are described in the original model by Kay and Hastings (41).

**Figure 1.**
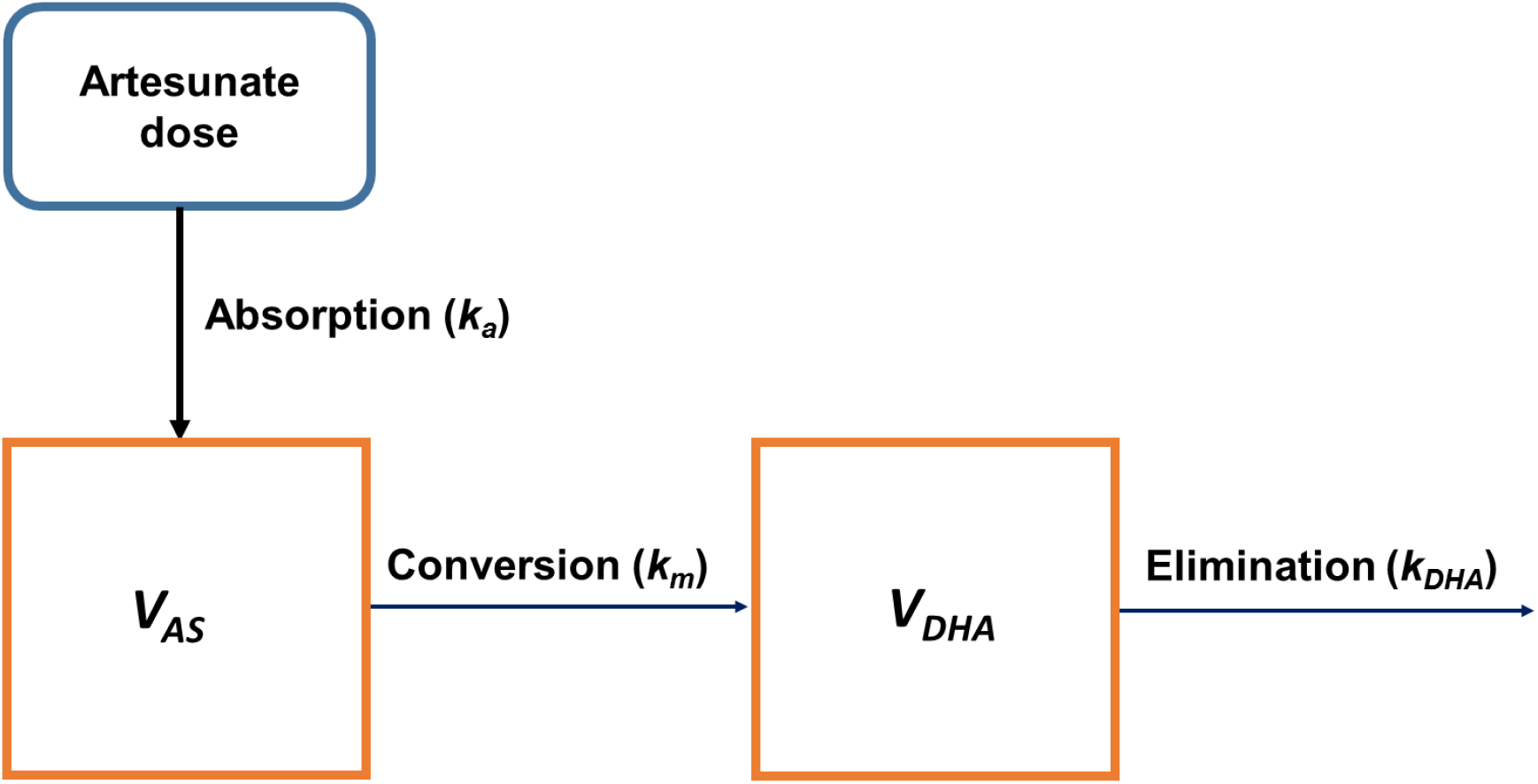
Structural pharmacokinetic model of artesunate (AS). AS follows a one-compartment disposition model, where it is absorbed from the gut at rate *k_a_* and converted into dihydroartemisinin (DHA) at rate *k_m_*. DHA is eliminated at rate *k_DHA_*. Note that full conversion of AS to DHA was assumed i.e. AS was not eliminated, only converted Abbreviations: AS: artesunate; DHA: dihydroartemisinin; *V_AS_*: volume of distribution of AS; *V_DHA_*; volume of distribution of DHA. This figure was adapted from Kay and Hasting (41).

For AQ and its active metabolite desethylamodiaquine (DEAQ), pharmacokinetics follow a more complex model. AQ and DEAQ were modelled as two separate drugs using a pair of two-compartment disposition models and a shared estimate of the absorption rate constant, *k_a_* (**Figure 2**); this approach was proposed and validated by Hietala et al (45). The model used the equation describing a drug with a three-compartment disposition (Equation 1.72 in (46)) but simplified to simulate two-compartments by setting the inter-compartmental clearance between compartments 1 (the central, blood compartment) and 3 (the unused peripheral compartment) to zero. The model equations assume the drug has first-order absorption, linear elimination and allows for multiple doses without lag time so that the amount of drug *C* present in the central compartment at time *t* is

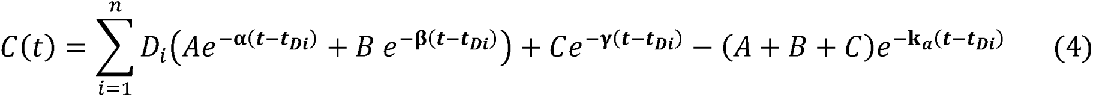

**Figure 2.**
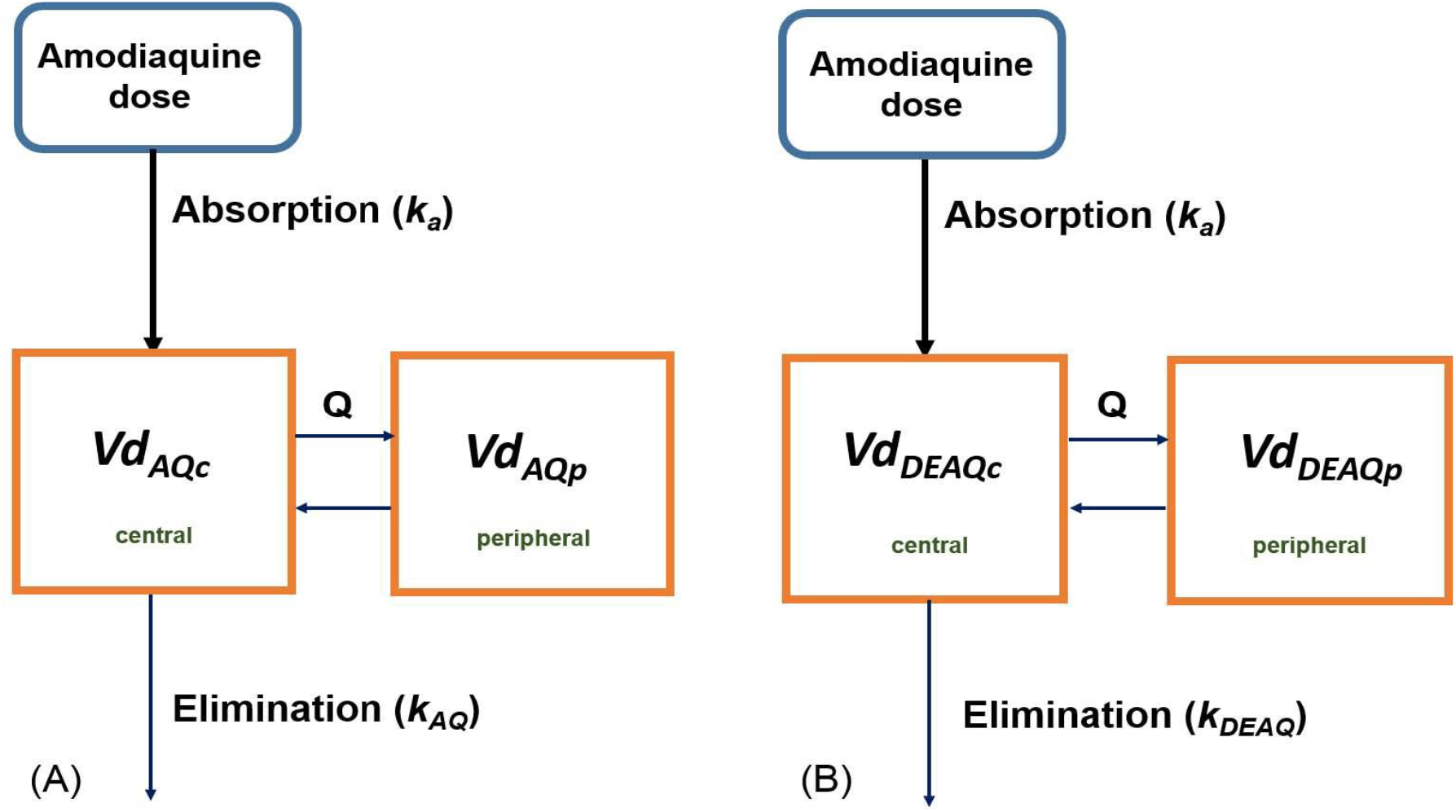
Structural pharmacokinetic model of amodiaquine (AQ) and desethylamodiaquine (DEAQ). The model includes two separate two-compartment models of AQ (Panel A) and its active metabolite DEAQ (Panel B). Both models share the same absorption rate (*k_a_*) but the remaining parameters were estimated independently for AQ and DEAQ. This figure was adapted from Hietala *et al.* (45). Abbreviations: *k_a_*: absorption rate constant; *Vd_c_*: central volume of distribution; *Vd_p_*: peripheral volume of distribution; AQ: amodiaquine; DEAQ: desethylamodiaquine; *k_AQ/DEAQ_*: elimination rate constants for AQ and DEAQ respectively; *Q*: distribution clearance between compartment.

where *D* is amount of drug (in mg) given in the *i*^th^ dose. A, B, C are macro-constants and α, β and γ are rate constants (see (46) for more details).

Both drugs in the AS-AQ combination have two active forms, an active parent drug and an active metabolite. The model therefore determines four drug concentrations and four corresponding estimates of drug dependent killing *f(C)* at each time step. The drug killing of the parent drug and its metabolite were assumed to have similar modes of action and so their effect cannot be additive (as implied in Equation 1). So at the end of each time step, only the drug form with the higher parasite-killing rate contributed to drug effect. The AS/DHA and AQ/DEAQ forms with the higher parasite-killing rate were combined assuming additive drug action to update parasite numbers (for more details see methods of (41)).

To simulate dosing according to age or weight, region-specific weight-for-age references for a global population and African, Asian, and Latin American populations were included as described previously (47). In brief, each individual was randomly assigned an age between six months to 25 years and their weight was read from the regional references (48) for a randomly selected weight-for-age percentile.

### Calibration and validation

Pharmacokinetic (PK) and pharmacodynamic (PD) parameters of AS and its metabolite DHA were previously validated in (41). The PK/PD parameters for AQ and DEAQ and their coefficient of variation (CV) estimates were extracted from the literature and are summarized in **Table 1**. For parameters where no CV was provided in the literature a default value of 30% was selected (as was done previously e.g. (40, 41)). It was assumed that parameters were normally distributed if the CV is ≤ 50% and log-normally distributed if they were > 50%.

**Table 1.** Antimalarial drug parameters for artesunate, amodiaquine and their active metabolites (DHA and DEAQ). The reported values are mean parameters (coefficient of variation).

| **Drug parameter** | **AS (41)** | **DHA (41)** | **AQ (45, 58)** | **DEAQ (45, 58)** |
| --- | --- | --- | --- | --- |
| $Vd_c$ (L/kg) | 46.6 (82) | 15 (48) | 8 (65) | 18.4 (48) |
| $Vd_p$ (L/kg) | – | – | 280 (26) | 68 (18) |
| $CL$ (L/hr) | – | – | 14 (22) | 0.66 (123) |
| $k_a$ (/day) | 23.98 (68) | – | 2.16 (30) | 2.16 (30) |
| $k_m$ (/day) | 11.97 (65) | – | – | – |
| $k$ (/day) | – | 44.15 (123) | – | – |
| $Q$ (L/hr) | – | – | 17 (26) | 1.3 (30) |
| $IC_{50}$ (mg/L) | 0.0023 (79) | 0.009 (117) | 0.0043 (163) (50) | 0.02 (132) (51) |
| $V_{max}^a$ (/day) | 27.6 | 27.6 | 2.3 (52) | 2.3 (52) |
| $n$ | 4 | 4 | 7 (53) | 7 (53) |

Abbreviations: $Vd_c$ : central volume of distribution; $Vd_p$ : peripheral volume of distribution; $CL$ : clearance; $k_a$ : absorption rate of parent drug; $k_m$ : conversion rate to active metabolite; $k$ : elimination rate of the drug or its metabolite; $Q$ : distribution clearance between compartments; $IC_{50}$ : 50% maximum inhibitory drug concentration; $V_{max}$ : maximal parasite-killing rate constant; $n$ : slope factor; AS: artesunate; DHA: dihydroartemisinin; AQ: amodiaquine; DEAQ: desethylamodiaquine.
<sup>a</sup> $V_{max} = -0.5 \times \ln(1/PRR)$ , where PRR is the parasite reduction ratio.

The PK parameters were validated by comparing the simulated maximal plasma concentration (*C_max_*) and time to *C_max_* (*T_max_*) of AQ and DEAQ to field observations (45). The PD parameters (40) were validated by matching simulated cure rates, parasite clearance times (PCT) and periods of chemoprophylaxis (PoC) to those estimated in clinical trials. The PCT is the time taken for the infection to fall below the limit of microscopic detection (defined here as <10^8^ parasites (49)). The PoC measures the time until new infections can occur after treatment, which is the duration of time that a drug suppresses the new infection. The PD parameters were taken from (50–53) and adjusted until the simulated cure rates, PCT and PoC matched field observations of the 28- day cure rate 96-98% (data from Nigerian children and Indian adults and children (13, 54) who were fully adherent and received age- or weight-based FDC regimens), the PCTs in these datatses were 1 ± 0.6 days (54–56) and PoC were 22 – 25 days (57). These results were obtained assuming full adherence to an age-based FDC or non-FDC regimens from clinical trials usually enrolled children from Africa. The final PK and PD parameters are given in Table 1 and are well within the range observed in the clinic and laboratory (41, 45, 50-53, 58).

### Simulation and analysis

This study compared the effectiveness and safety of different treatment regimens and drug formulations for the adherence scenarios listed in **Table 2**. Weight-based dosing regimens with fixed-dose and co-blistered combinations of AS-AQ were based on drug dose per patients’ weight band as recommended by WHO ((2); **Table 3**). Note, individuals under 5 kg were excluded from this analysis and replaced by a new random sample record. Age-based dosing regimens were selected from clinical trials (11, 59, 60) as WHO recommends only weight-based dosing regimens. The age- and weight-based regimens used in this study are presented in **Table 3**.

**Table 2.** Simulated scenarios.

| **Scenario** | **AS treatment schedule** | **AQ treatment schedule** |
| --- | --- | --- |
| 1 Baseline (fully adherent) | Once daily for three days | Once daily for three days |
| 2 Omitting the third dose | Once daily for two days | Once daily for two days |
| 3 Omitting the second and third dose | Single dose | Single dose |
| 4 Delaying doses for 12 hours | At 0, 36, and 60 hours | At 0, 36, and 60 hours |
| 5 AQ monotherapy | – | Once daily for three days |
| 6 AS monotherapy | Once daily for three days | – |

Abbreviations: AQ: amodiaquine; AS: artesunate.

**Table 3.** Simulated dosing regimens.

|  | **Regimen** | **AS dose** | **AQ dose** |
| --- | --- | --- | --- |
| A | WHO target dose (2) | 4 mg/kg | 10 mg/kg |
|  | **Weight-based** |  |  |
| B | Study 1 – FDC (2) |  |  |
|  | 4.5-8.9 kg | 25 mg | 67.5 mg |
|  | 9-17.9 kg | 50 mg | 135 mg |
|  | 18-35.9 kg | 100 mg | 270 mg |
|  | > 36 kg | 200 mg | 540 mg |
| C | Study 1 – co-blistered (2) |  |  |
|  | 4.5-8.9 kg | 25 mg | 76.5 mg |
|  | 9-17.9 kg | 50 mg | 153 mg |
|  | 18-35.9 kg | 100 mg | 306 mg |
|  | > 36 kg | 200 mg | 612 mg |
|  | **Age-based** |  |  |
| D | Study 2 – FDC (11) |  |  |
|  | 6-11 month | 25 mg | 67.5 mg |
|  | 1-5 years | 50 mg | 135 mg |
|  | 6-13 years | 100 mg | 270 mg |
|  | ≥ 14 years | 200 mg | 540 mg |
| E | Study 3 – co-blistered (59) |  |  |
|  | < 1 year | 25 mg | 76.5 mg |
|  | 1-5 years | 50 mg | 153 mg |
|  | 6-12 years | 100 mg | 306 mg |
|  | >13years | 200 mg | 612 mg |
| F | Study 4 – FDC (60) |  |  |
|  | 1-5 years | 50 mg | 135 mg |
|  | 6-13 years | 100 mg | 270 mg |
|  | ≥ 14 years | 200 mg | 540 mg |
| G | Study 4 – co-blistered (60) |  |  |
|  | 1-5 years | 50 mg | 153 mg |
|  | 6-10 years | 100 mg | 306 mg |
|  | 11-15 years | 150 mg | 459 mg |

Abbreviations: WHO: World Health Organization; AS: artesunate; AQ: amodiaquine; FDC: fixed-dose combination

The details of the different model runs are outlined in **Table 4**. A single model ‘run’ simulates 10,000 individuals for each of the regional weight-for-age distributions (representative of either a global, African, Asia or Latin American populations) and followed patients for 28 days after the start of treatment (i.e. the recommended follow-up duration according to the WHO guidelines (26)). Treatment outcome at 28 days was defined as follows; 1) all parasites cleared (*P_t_* < 1); 2) parasite are still present but below limit of microscopic detection (LoD) of *P_t_* <10^8^ and might subsequently either clear or recrudesce; 3) recrudescence which fell below LoD at some point during follow-up but later recrudesced to above LoD at some point before the end of the 28 day follow-up; 4) parasites are never cleared (*P_t_* >10^8^) and are always detectable during entire post-treatment period. Categories 1) and 2) were classified as clinical cures and categories 3) and 4) were classified as clinical failures (this corresponds to what is defined as cured/failures in the field).

**Table 4.**
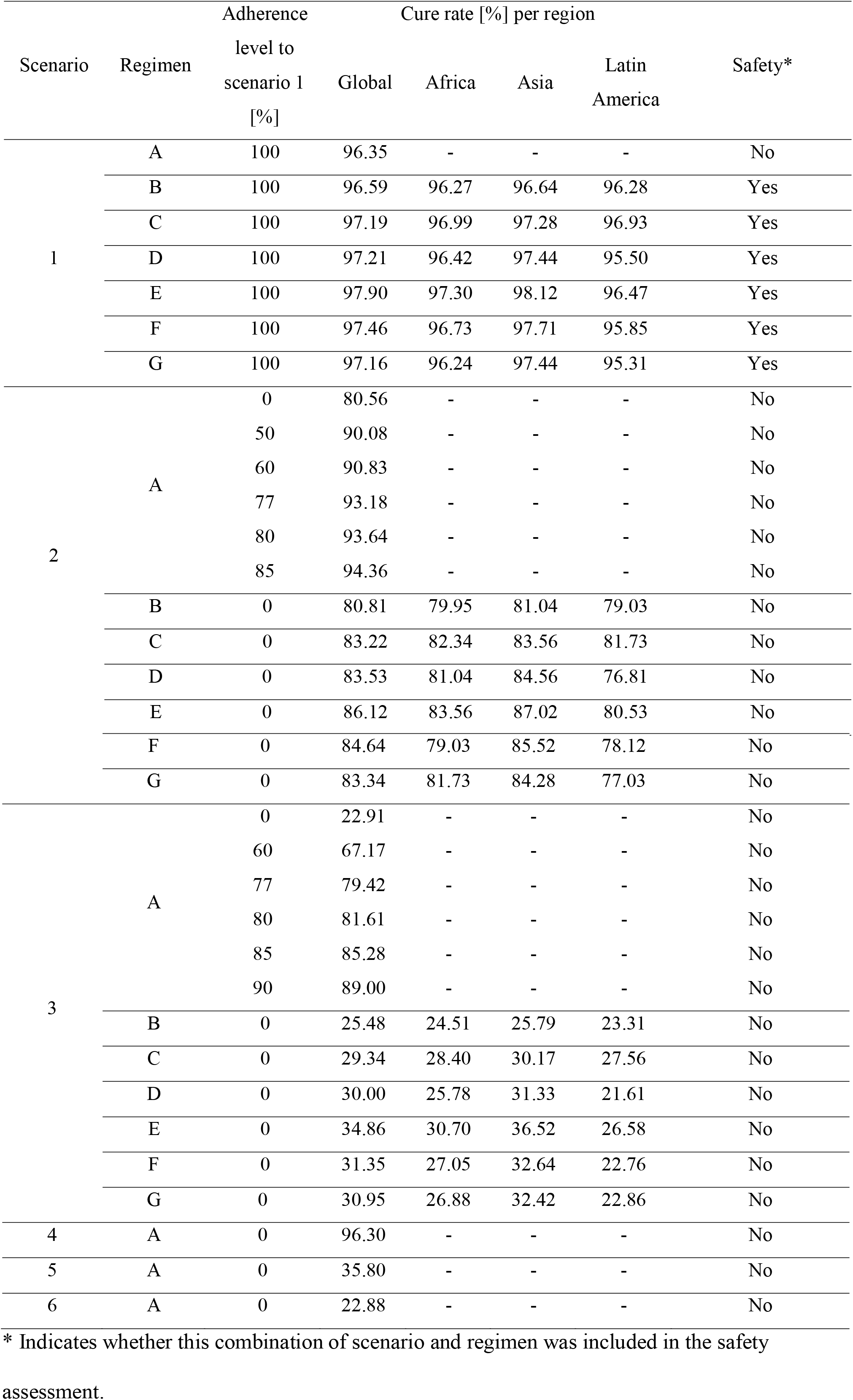
Cure rates for different combinations of dosing regimens, adherence scenarios and adherence levels. Details of scenarios and regimens can be found in Table 2 and Table 3 respectively. Follow-up time was 28 days. The adherence level column indicates the proportion of simulated individuals (n=10,000) following the recommended dosing for each regimen according to scenario 1 (i.e. once daily dosing for three days). For example, when simulating scenario 2 with regimen A at an adherence level of 77%, means 77% of patients (n=7,700) followed scenario 1 with regimen A, and the remaining 23% of patients (n=2,300) missed the last dose of treatment according to scenario 2 with regimen A.

To reproduce results of drug effectiveness in a real life setting, the model accounted for missed doses and poor adherence, using patterns of non-adherence found in the literature (61, 62). An adherence level defined the proportion of patients that followed the recommended treatment regimens; seven adherence levels were investigated here i.e. 50%, 60%, 77%, 80%, 85%, 90% and 100%. So an adherence level of 77% for example, means 77% of patients followed the full three day course of AS-AQ treatment and 23% followed one of the alternative scenarios listed in **Table 2**.

Safety was assessed by determining the proportion of patients who received an AS and AQ dose within the therapeutic range. The therapeutic range of AS was defined as 2–10 mg/kg/day and 7.5 –15 mg/kg/day for AQ (2). The magnitude of under- and overdosing of AS and AQ was determined as well as median values of *C_max_* of AS, DHA, AQ, and DEAQ.

## Results

### Efficacy and effectiveness of the WHO recommended regimen

Efficacy describes how a drug performs under ideal conditions for example, when treatment is directly observed in clinical trials. In contrast, effectiveness describes how well a drug works in a real-life setting where, for instance, patients take medication unsupervised. The efficacy of the WHO recommended target dose of 4 mg/kg AS and 10 mg/kg AQ daily (regimen A) with full-adherence (scenario 1) was compared with the effectiveness of the same regimen in patients with varying degrees of non-adherence (scenarios 2–6) at day 28 of follow-up (**Table 4** and **Figure 3**). Scenario 1 assumed all individuals took the full three days of treatment and resulted in a cure rate of approximately 96%. When the third dose was missed (scenario 2) cure rates decreased to 80–94% depending on the proportion of non-adherent patients (**Figure 3A**). However, the WHO regimen appears relatively robust with clinical cure rates remaining above 90% when half the patient population only takes the first two treatment doses. In contrast, when 10% of patients omitted both the second and third doses (scenario 3) cure rates dropped to 89% (**Figure 3B**). The proportion of patients cured drops to 23% when only the first dose of treatment is taken. Delaying the second dose by 12 hours (scenario 4) did not alter the drug effectiveness. Monotherapies of either AQ or AS given for three days (scenarios 5 and 6 respectively) showed low cure rates of 36% and 23% respectively (**Table 4**).

**Figure 3.**
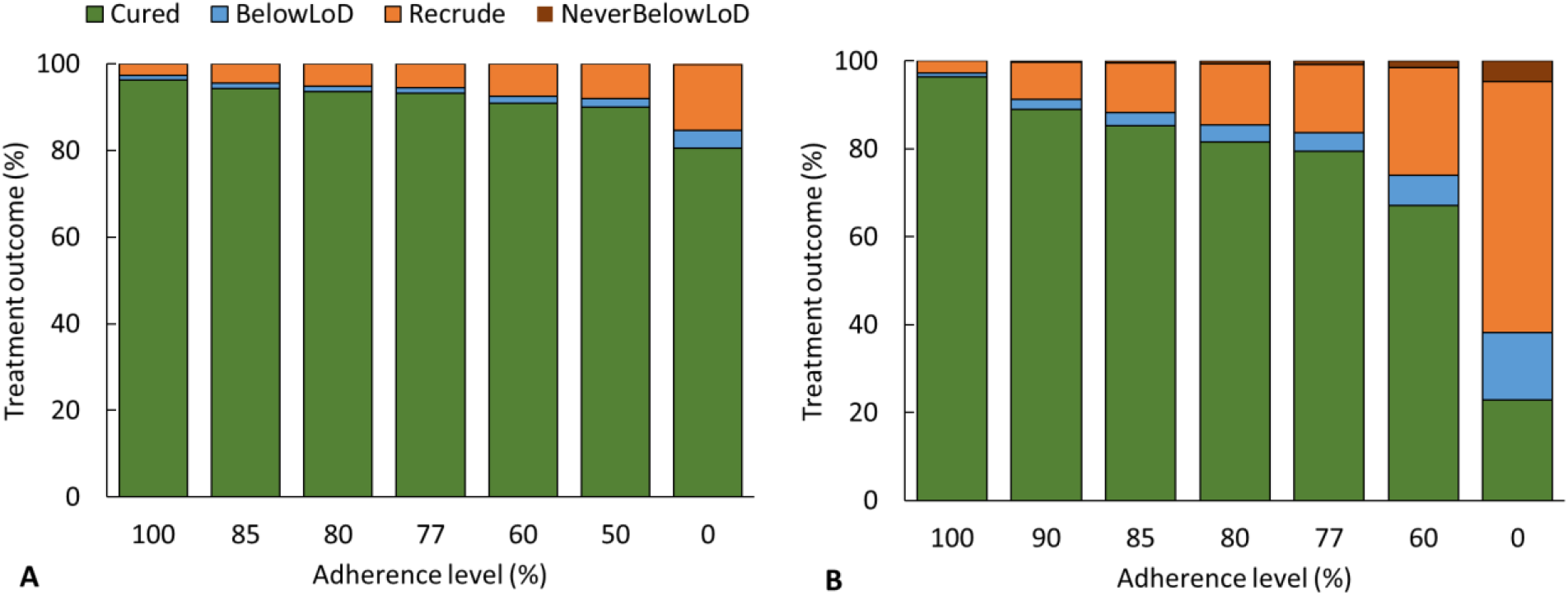
The impact of patient adherence on treatment outcome. This was in patients prescribed regimen A (7), i.e. prescribed the therapeutic target dose of 4 mg/kg/day for artesunate and 10 mg/kg/day for amodiaquine (26) (**Table 3**). Each bar in the plots represents a decreasing proportion of fully-adherent patients in the simulated population. Non-adherent patients were assumed to take their regimen according to **(A)** Scenario 2, i.e. missed their third treatment dose (**Table 2**), or **(B)** Scenario 3, i.e. missed both their second and third doses (**Table 2**). Abbreviation: LoD: limit of microscopic detection; Recrude: Recrudescence

In order to measure the immediate therapeutic response to the drugs, PCT was calculated. In scenarios 1 and 4, the median PCT was 1.25 days and the median time to recrudescence was 22 days (data pooled over all the regimens) in both cases (**Figure 4**). The median PCT did not change until patients missed more than one dose of the combination. As expected, monotherapies with the fast acting, short-lived component AS, had a negligible impact on the median PCT but shortened the median time to recrudescence. Conversely, monotherapies with the slower acting, longer-lived AQ partner drug lengthened the median PCT but did not affect the time to recrudescence.

**Figure 4.**
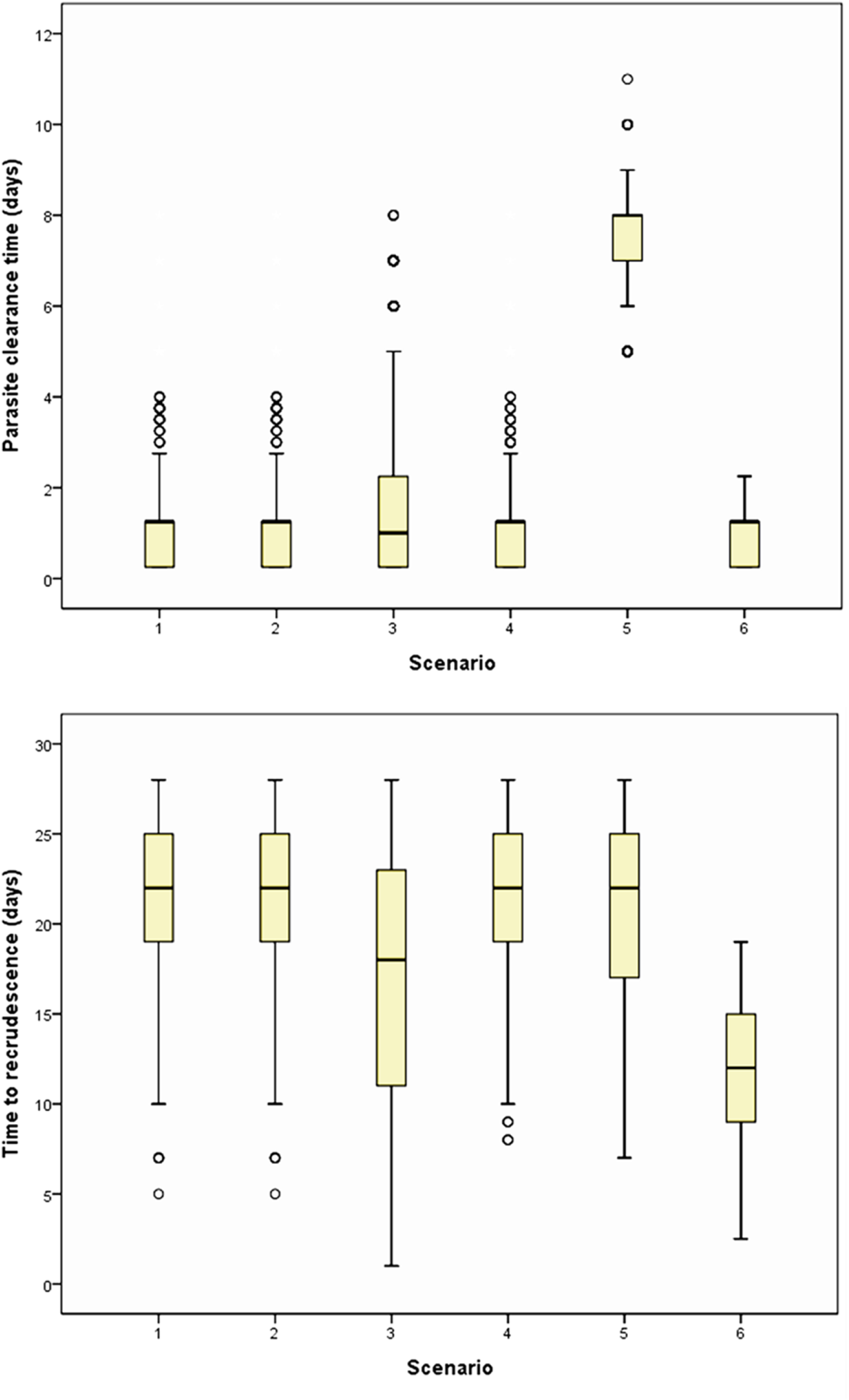
Distribution of parasite clearance time (PCT) and time to recrudescence (TTR) (**28**). PCT and TTR following treatment with artesunate-amodiaquine (7) and different scenarios (detailed in **Table 2**).

### Efficacy and effectiveness

The cure rates of the six simulated age- and weight-based regimens (B to G, **Table 3**) were all above 96% when patients were fully adherent to the three day regimen (scenario 1, **Table 4**). Cure rates in the three regions were compared for each regimen and, were highest in Asia (97%) and lowest in Latin America (95%) assuming complete adherence to the regimen (**Figure 5A**). When all patients followed scenario 2 (i.e. omitted the third doses), the cure rates dropped below 90% in all six regimens (**Figure 5B**) and when all patients followed scenario 3 (i.e. omitted the second and third doses), the cure rates dropped to below 35% (**Figure 5C**). These results emphasize that full three day course of AS-AQ is necessary to treat uncomplicated falciparum malaria.

**Figure 5.**
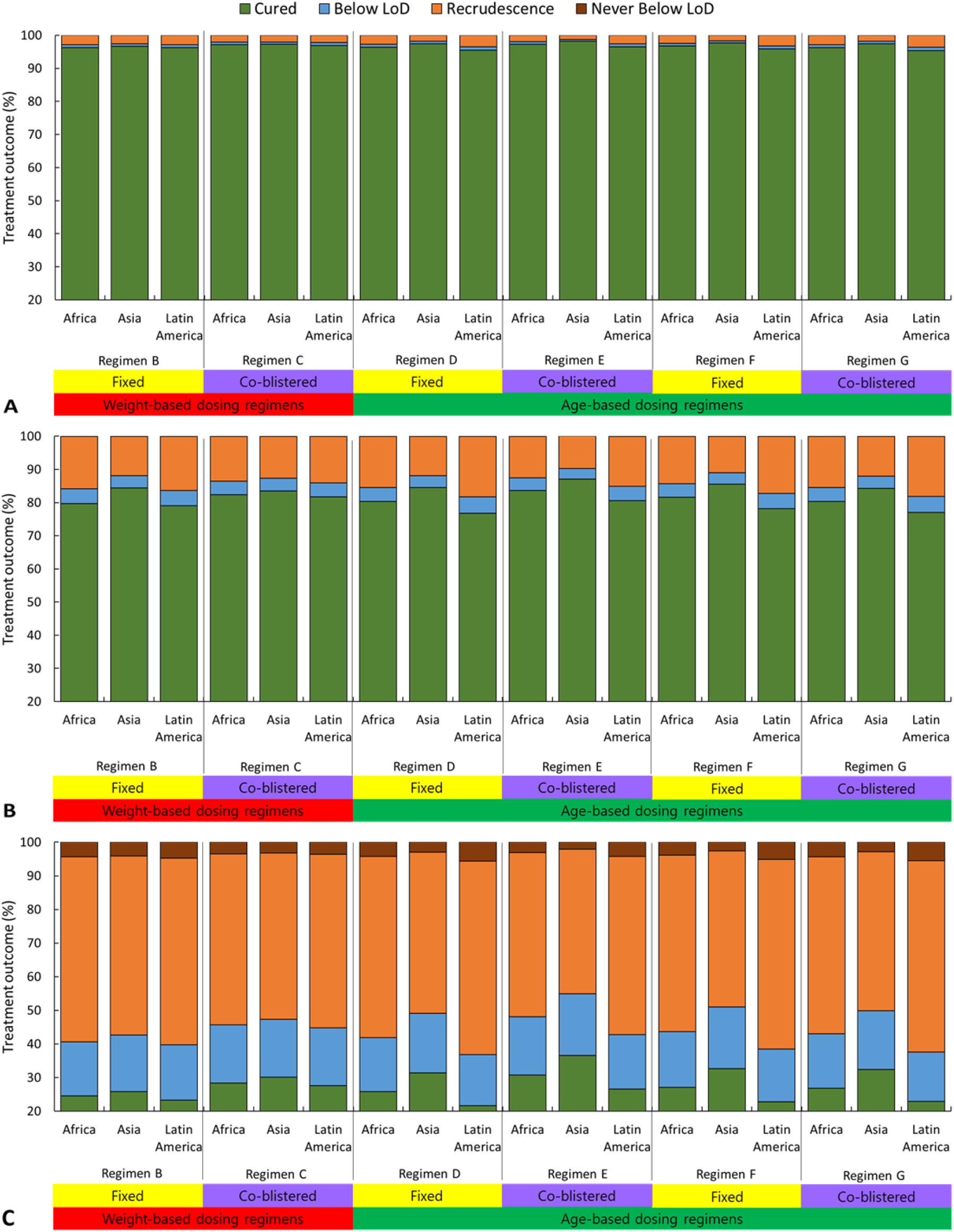
Treatment outcome in patients receiving age- or weight-based dosing regimens of artesunate-amodiaquine. Scenario 1 treatment outcome for regimens B to G (**Table 3**) simulated using three different region-specific weight-for-age references (Africa, Asia and Latin America). The panels represent the patients **(A)** were fully adherent (Scenario 1); **(B)** omitted one dose of AS-AQ (Scenario 2); **(C)** omitted two doses of AS-AQ (Scenario 3)

### Drug dosing and safety evaluation

In scenario 1, all regimens showed high cure rates (i.e. >95 %) but selection of the ‘best’ or optimal regimen should also depend on the regimen’s safety profile. In absence of an absolute ‘toxic’ mg/kg dose or plasma concentration threshold, relatively ‘safe treatments’ were determined in the simulations using the proportion of patients who received doses in the recommended therapeutic range of AS and AQ. The therapeutic range was defined as 2–10 mg/kg/day for AS and 7.5–15 mg/kg/day for AQ (26). **Table 5** and **Figure 6** show the proportion of patients under- or overdosed with either AS or AQ for each of the regimens described in **Table 3**. In the simulated populations 15% of patients received doses above and 4.4% below the therapeutic dose range of AQ. This was an average across all regimens and regions but there was considerable differences between regimens. For example, no patients following Regimen B (in all three continents) were overdosed. In addition, the two age-based dosing regimens seems more likely to over- or under-dose AQ. Asians are more likely to be overdosed and Latin Americans more likely to be under dose than any other regions (**Figure 6**).

**Table 5.** Dosing accuracy and predicted exposure depending on dosing regimen. This table shows the percentage of patients from different regions receiving doses outside the therapeutic range when given one of the six regimens described in **Figure 3**, the drug dosage and predicted maximal plasma concentration of drug and its metabolites. Values of dosage and *C_max_* are in medians.

| | OD AS<br>[%] | UD AS<br>[%] | OD AQ<br>[%] | UD AQ<br>[%] | AS dose<br>[mg/kg/<br>day] | AQ dose<br>[mg/kg/<br>day] | $C_{max}$ AS<br>[mg/dL] | $C_{max}$ DHA<br>[mg/dL] | $C_{max}$ AQ<br>[mg/dL] | $C_{max}$ DEAQ<br>[mg/dL] |
| --- | --- | --- | --- | --- | --- | --- | --- | --- | --- | --- |
| **Regimen A** |  |  |  |  |  |  |  |  |  |  |
| Africa | 0.0 | 0.0 | 0.0 | 0.0 | 4 | 10 | 0.5 | 0.8 | < 0.1 | 0.3 |
| Asia | 0.0 | 0.0 | 0.0 | 0.0 | 4 | 10 | 0.5 | 0.8 | < 0.1 | 0.3 |
| Latin America | 0.0 | 0.0 | 0.0 | 0.0 | 4 | 10 | 0.5 | 0.8 | < 0.1 | 0.3 |
| **Regimen B** |  |  |  |  |  |  |  |  |  |  |
| Africa | 0.0 | 0.0 | 0.0 | 1.7 | 3.3 | 8.9 | 0.5 | 0.7 | < 0.1 | 0.3 |
| Asia | 0.0 | 0.0 | 0.0 | 0.0 | 4.2 | 11.3 | 0.5 | 0.8 | < 0.1 | 0.4 |
| Latin America | 0.0 | 0.0 | 0.0 | 3.2 | 3.8 | 10.1 | 0.5 | 0.8 | < 0.1 | 0.4 |
| **Regimen C** |  |  |  |  |  |  |  |  |  |  |
| Africa | 0.0 | 0.0 | 13.8 | 0.4 | 3.9 | 11.8 | 0.5 | 0.8 | 0.1 | 0.4 |
| Asia | 0.0 | 0.0 | 23.2 | 0.0 | 4.2 | 12.8 | 0.5 | 0.8 | 0.1 | 0.4 |
| Latin America | 0.0 | 0.0 | 11.6 | 0.7 | 3.8 | 11.5 | 0.5 | 0.8 | < 0.1 | 0.4 |
| **Regimen D** |  |  |  |  |  |  |  |  |  |  |
| Africa | 0.0 | 0.1 | 9.1 | 6.0 | 4.0 | 10.7 | 0.5 | 0.8 | < 0.1 | 0.4 |
| Asia | 0.0 | 0.0 | 24.0 | 1.4 | 4.8 | 13.0 | 0.6 | 1.0 | 0.1 | 0.4 |
| Latin America | 0.0 | 1.7 | 2.5 | 16.1 | 3.6 | 9.6 | 0.5 | 0.7 | < 0.1 | 0.3 |
| **Regimen E** |  |  |  |  |  |  |  |  |  |  |
| Africa | 0.0 | 0.0 | 23.8 | 1.2 | 4.1 | 12.4 | 0.5 | 0.8 | 0.1 | 0.4 |
| Asia | 0.1 | 0.0 | 50.9 | 0.3 | 4.9 | 15.0 | 0.6 | 1.0 | 0.1 | 0.5 |
| Latin America | 0.0 | 0.9 | 9.8 | 5.0 | 3.6 | 11.1 | 0.5 | 0.7 | < 0.1 | 0.4 |
| **Regimen F** |  |  |  |  |  |  |  |  |  |  |
| Africa | 0.1 | 0.0 | 13.6 | 4.1 | 4.1 | 11.1 | 0.5 | 0.8 | < 0.1 | 0.4 |
| Asia | 0.2 | 0.0 | 29.8 | 0.7 | 5.0 | 13.4 | 0.6 | 1.0 | 0.1 | 0.5 |
| Latin America | 0.0 | 1.1 | 5.2 | 12.5 | 3.7 | 10.0 | 0.5 | 0.7 | < 0.1 | 0.3 |
| **Regimen G** |  |  |  |  |  |  |  |  |  |  |
| Africa | 0.1 | 1.2 | 19.4 | 9.1 | 3.6 | 10.9 | 0.5 | 0.7 | < 0.1 | 0.4 |
| Asia | 0.2 | 0.0 | 32.2 | 0.9 | 4.2 | 12.8 | 0.6 | 0.8 | 0.1 | 0.5 |
| Latin America | 0.0 | 3.0 | 8.1 | 16.6 | 3.1 | 9.5 | 0.4 | 0.6 | < 0.1 | 0.3 |

Abbreviations: AS: artesunate; AQ: amodiaquine; DHA: dihydroartemisinin; DEAQ: desethylamodiaquine; $C_{max}$ : maximal concentration of drug; OD: over-dosed; UD: under-dose

**Figure 6.**
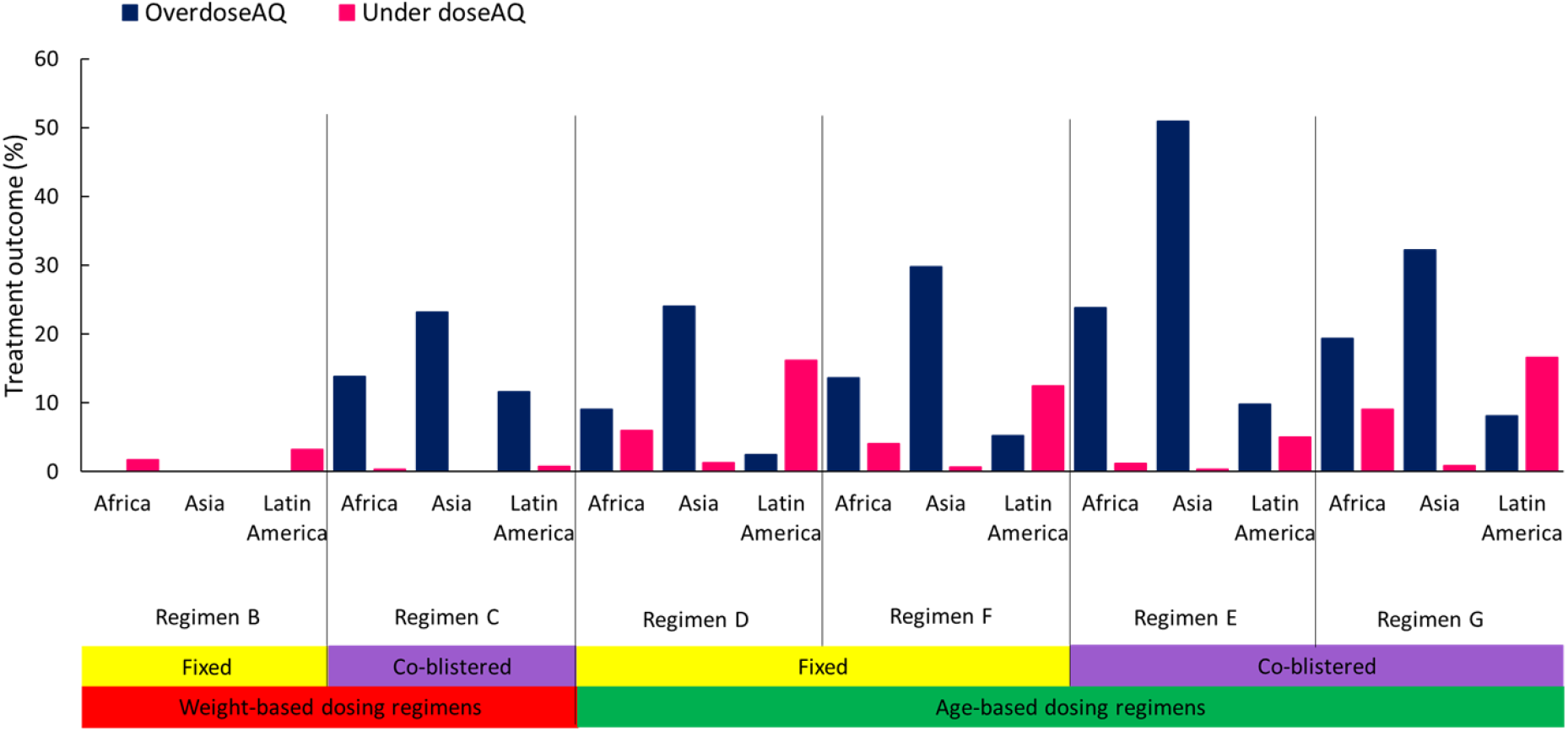
The percentage of patients over- or under-dosed with amodiaquine (AQ) depending on regional weight-for-age distribution. Patients were dosed according to either weight (red) or age (green) and given either a fixed dose formulation (yellow) or a co-blistered formulation (purple) of artesunate-amodiaquine. Regimens B to G were as described in **Table 3**.

The specific dose of AQ each patient received is given on **Figure 7** using examples of one age- and one or weight-based regimen (**Table 3**, regimens B and D respectively). The weight-based dosing regimen with FDC (regimen B) showed that overdosing of AQ was uncommon in all ages (**Figure 7A**). The age-based dosing regimen showed that patients near the lower cutoff of each age band tended to receive higher mg/kg dosages and so were more frequently overdosed with AQ (**Figure 7B**). The proportion of patients overdosed decreased with increasing patient age with a small proportion of patients at the upper dosing band cut-offs being under-dosed (**Figure 7B**).

**Figure 7.**
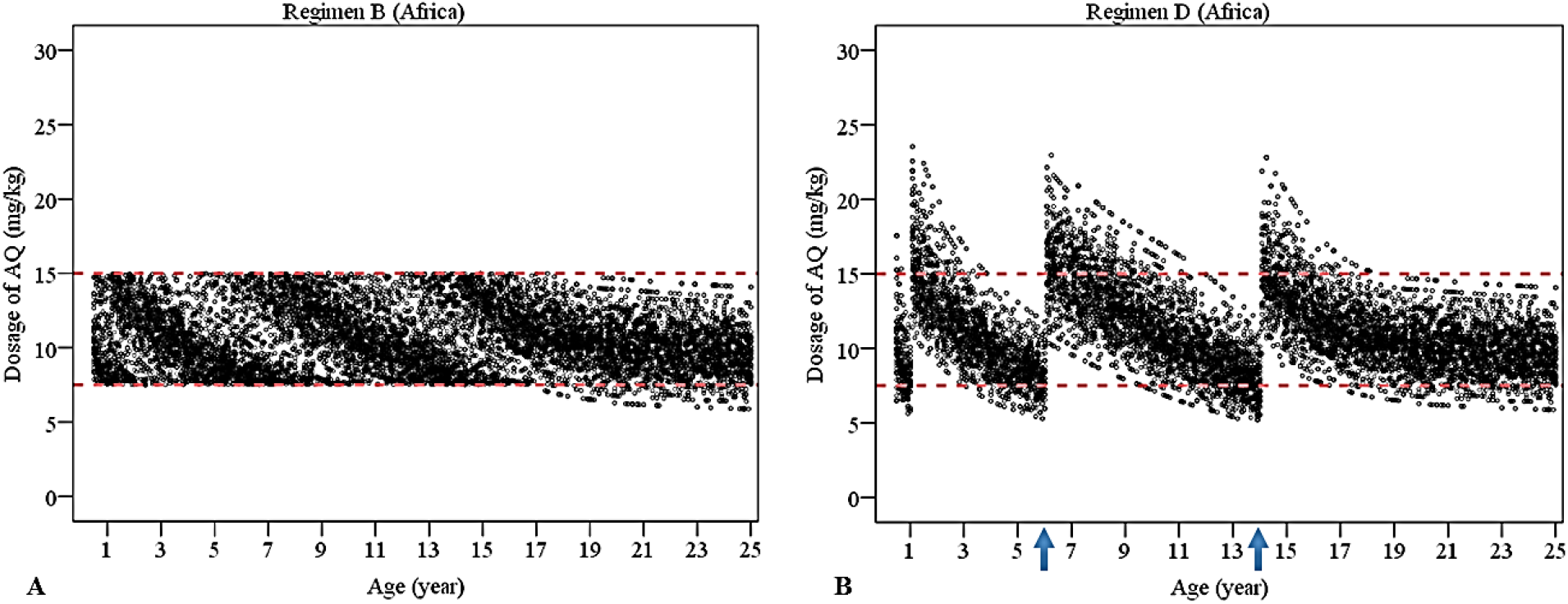
Amodiaquine (AQ) dosage received by simulated patients dosed according to weight or age. Simulated regimens are listed in **Table 3. (A)** Weight-based regimens (regimens B). **(B)** Age-based regimens (regimens D). The red dotted lines indicate the therapeutic dose range of AQ (26). The blue arrows indicate the cut-off points of the age-based regimen.

## Discussion

This study adapted an existing antimalarial pharmacological model to simulate AS-AQ treatment and compare the efficacy, effectiveness and safety of a variety of AS-AQ regimens based on fixed dose and non-fixed formulations and using either age- or weight-based dosing bands. The structural model used herein for AQ and its active metabolite was described previously (45), but the literature provides several alternative structural PK models (45, 63, 64). AQ and DEAQ were modelled as separate drugs with two-compartment PK model structures because it can be assumed that AQ and DEAQ have similar physiochemical characteristics such as their absorption, distribution, solubility and stability. Furthermore, the separate two-compartment PK models have an algebraic solution (41, 46) that makes it computationally preferable to a linked four-compartment PK model. Recently, more sophisticated PK models have become available using non-linear mixed-effect modeling (65) and our analysis could be extended to incorporate these more complex dynamics. In the meantime, we note that the simpler models provide a good fit to PK data and therefore constitute a good platform for our primary objectives of investigating the effects of age- and weight-based dosing bands and the impact of poor patient adherence.

The literature used to calibrate the PK model for AQ reported large inter-individual variation of PK parameters (45, 66), i.e. for absorption rate the CV was 100% and for volume of distribution the CV was 139%. Despite this, those CVs originating from a study that enrolled children were retained because they came from the study with the largest sample size. The CVs for the other PK parameters, i.e. *k_a_, CL, Vd_c_*, were too narrow and resulted in model output that was inconsistent with field data (45, 53). Consequently, CVs from a separate study that enrolled women with *P. vivax* malaria during and after pregnancy were used (58). Although, the volume of distribution in pregnant women was larger than in non-pregnant women, the variation of the PK parameters in that study was smaller than the one involving children. Where no CV for AQ/DEAQ specific PK and PD parameters could be found in the literature, the CV was set to 0.3 (for further discussion see (41)). Note that the model ignored the acquired immunity of malaria as there is currently no consensus mathematical description of immune acquisition (see for example, (40, 67-69)) but it is important that the model can predict efficacy and effectiveness in the most vulnerable, i.e. non-immune, individuals. The cure rate for validation was thus selected from studies conducted in Asia or Africa (the latter with pediatric patients) because these populations are presumed to have relatively low immunity to falciparum malaria. Overall, the pharmacological model used here was able reproduce field data (13, 45, 54, 56, 70).

The model generated cure rates, recrudescence rates and the safety profiles for six different regimens. The efficacies of all six regimens were high when patients adhered to the full three-day course of AS-AQ (scenario 1 and 4) and, as expected, lower when patients did not follow the three-day course (scenario 2 and 3). Monotherapies of either drug taken for three days (scenario 5 and 6) also showed low cure rates as reported from clinical trials comparing monotherapy to combination therapy (13, 55). This study also investigated safety of different dosing regimens and combinations. The weight-based dosing with FDC (regimen B) was the best option in regard to safety and was similar in terms of efficacy compared to the other regimens. Over- and under-dosing of AQ was a significant problem in the age-based dosing regimens.

The pharmacological model generated similar proportions of cure rates with those observed in clinical trials for AS-AQ three day course treatments. Many clinical trials reported polymerase chain reaction (PCR)-corrected cure rates of 97% or above at day 28 or 42 after treatment when patients were treated with 10mg/kg of AQ for three days combined with artesunate (11, 13, 60, 71). The model also generated 97% cure rate for full adherence. However, the model cure rates following AQ monotherapy (10mg/kg once daily for three days) were just 36% if cure rate is defined as patients who clear all parasites or 68% if cure rate is defined as patients who had parasites below the limit of microscopic detection. This is much lower than the day 28 PCR-corrected cure rates from equivalent clinical trials using AQ monotherapy, i.e. 88% in India (13), 54% in Kenya (72), 79% in Senegal (72) and 85% in Gabon (72). In addition, when model follow-up to 28 days, the simulated cure rates dropped to 36% (defined as the proportion of patients with undetectable parasites). The IC_50_ values for AQ and DEAQ in this study might reflect resistant strains, thereby generating a lower cure rate than clinical trials. When IC_50_ was decreased from 0.02 mg/L to 0.01 mg/L, model cure rates become more similar to those reported in the studies i.e. 54% of patients cleared all parasites and 84 % of patients had undetectable parasites. These result suggests that *P. falciparum* is likely to be sensitive to AQ in Gabon but not in Kenya(72–74); possible reflecting the much higher usage of AQ in West-compared to East-Africa. As expected, these results indicate that monotherapy had risks of treatment failure and emergence of drug resistance. Therefore, intensive effort is required to reduce monotherapy treatment by improving patients’ adherence.

Adherence to ACT varies across regions, ranging from 48-94% (23, 75). To increase patient adherence, AS-AQ treatment is now available in co-blistered packaging or as FDC and the reported efficacies are more than 95% (29, 60). The Worldwide Antimalarial Resistance Network (WWARN) recently conducted a pooled analysis that included 43 studies with 9,106 patients treated with a three day course of AS-AQ for uncomplicated falciparum malaria (7). Their study compared the efficacy of several AS-AQ combinations and found the efficacy of FDC (98.1%) and co-blistered non-FDC (97.9%) were similarly high but found the efficacy of loose non-FDC-30 (95%) was statistically significantly lower (7). The median AQ dosed received in the WWARN’s pooled analysis was 32.4 mg/kg for FDC (AS/AQ ratio 2.7) and 35.3 mg/kg for co-blister non-FDC (AS/AQ ratio 3.06) (7), which was very similar to that used in this model, i.e. a median AQ dose of 32.8 mg/kg for FDC and 35.9 mg/kg in co-blister non-FDC, and resulted in very similar AS/AQ ratios. It is gratifying to note the simulated cure rates for AS-AQ treatment with FDC and co-blistered non-FDC were highly consistent to the findings from the WWARN study. Patients in both the WWARN pooled analysis and the simulation receiving non-FDC formulations, tended to have slightly higher median dose of AQ. The proportion of patients receiving AQ doses below the therapeutic dose range was similarly consistent at 3.4% and 4.4% in the analysis by the WWARN AS-AQ study group and the simulation respectively. The only small deviation between the results of WWARN and the simulated patients occurred when measuring the proportion of patients under-dosed. WWARN report under-dosing with FDC in 1.1% of patients and with co-blister non-FDC in 0.9% (7) of patients while the simulation predicted under-dosing in 5.1% and 3.8% respectively. Since FDC contains less amount of AQ than co-blister combination, FDC had higher risk of under-dosing. However, the difference was small. Under-dosing has the potential to affect efficacy and effectiveness and the model predicted results highly consistent with those presented in WWARN’s pooled analysis (7) except when simulating the proportion of patients receiving sub-therapeutic AQ doses.

Patients overdosed with AQ are more prone to vomiting than those receiving the appropriate dose (76), a side effect that can make patients and caregivers reluctant to administer treatment (61). When treating children with AS-AQ, overdosing of the AQ component is a particular concern regardless of whether children are dosed according to their age or weight. A study in Senegal (10) gave patients co-blistered AS-AQ (AS 50 mg, AQ 153 mg) based on weight or age; 2.1% of patients were overdosed with AQ when dosed according to weight and 17.8% were overdosed with AQ when dosed according to age. WWARN’s pooled AS-AQ analysis (7) showed children received the lowest AQ total dose if they were under 12 months old (28.9 mg/kg FDC vs 32.6 mg/kg co-blistered non-FDC) and the highest AQ per dose if they were 1–4 years old (33.8 mg/kg FDC vs 38.3 mg/kg co-blistered non-FDC) (7). The simulated results presented here show similar patterns of under-dosing: Patients less than 12 months old received 28.2 mg/kg with FDC and 32.0 mg/kg with non-FDC and children aged 1–4 years old received 35.7 mg/kg with FDC and 40.5 mg/kg with non-FDC. Non-FDC formulations also tended to result in slightly higher amounts of AQ in all age groups and this trend was similar in the WWARN study (7). The authors of the WWARN study credited higher failure rates for the non-FDC products to the fact that the FDC product “was developed using a weight-for-age reference database from malaria endemic countries, to ensure optimal dosing with the pediatric formulation”. When “applied either by weight- or age-based criteria” the FDC product “probably increases dosing accuracy, and the availability of different tablet strengths, including a pediatric formulation, obviates the need for tablet splitting, reduces the pill burden and potentially improves adherence”

The WHO recommends that first-line treatment should be changed when failure rates exceed 10% (2). In this study, the clinical failure rate was less than 5% (assuming full adherence for all regimens) indicating that current treatment policies are adequate. Clinical failure rates increased dramatically to 15% when patients missed one dose of AS-AQ. As per WHO recommendations failure rate of 15% are expected to result in a change of treatment regimen. However, the importance of ensuring patient adherence with recommended drug doses and schedules was also simulated. Non-adherence was incorporated in the models and used to show that cure rates could be improved to above 90% if less than half the simulated population missed one dose of AS-AQ combination treatment or if no more than 10% of the population missed two doses of AS-AQ. Clearly careful monitoring of patient adherence is needed for successful control of malaria but is difficult to quantify in practice (23). Non-adherence often leads to more patients with sub-therapeutic drug levels. This causes two main problems, first more patients fail treatment and often require re-treating which in turn results in higher treatment costs and second, sub-therapeutic drug levels may increase the appearance and subsequent spread of drug resistance.

The WHO guidelines (2) recommend that treatment doses be calculated according to the patients’ body weight (30). In most of the malaria endemic countries, dosing by age is a more common practice, in some cases due to the lack of functional scales (77) but more often because health workers prefer to prescribe according to age for simplicity (31). This study showed that simulated patients who received age-based doses had highest AQ levels in the younger ages within each age band (not surprising given that younger patients tend to be lighter). A clinical trial in Senegal showed that dosing patients according to body weight resulted in more correct AQ doses; 18% of patients were overdosed with AQ when treated according to age and 13% were overdosed with AQ when treated according to body weight (10). These results show the benefits to patients that result from the use of regional weight-for-age distributions to optimize age-based dosing regimens (78).

When comparing the results across different populations globally, these simulated results show Asian populations are at the highest risk of receiving AQ above the therapeutic range and Latin American populations had the highest risk of receiving AQ below the therapeutic range. This become particularly apparent when simulated patients were dosed according to their age. One possible explanation is that Asian people tend to weigh less than Latin Americans (48). This reinforces the importance of using regional weight-for-age distributions to inform treatment decisions at a regional or country level.

## Conclusion

In conclusion, the simulated probability of receiving an AQ over-dose and predicted drug effectiveness were highly consistent with clinical trials. Efficacy and effectiveness were high in all regimens when simulating fully adherent patients, but FDCs have a major advantage in ensure patients take both dugs in the ACT so minimizing the risks associated with patients taking AS or AQ monotherapies. The FDC weight-based dosing regimens had better proportion of patients receiving the recommended target dose and will remain the preferred regimen. However, it must be recognized that this is not always feasible, whether it is due to technical difficulties locally, healthcare preference or simply a logistical problem, for example in large MDA campaigns. Age-based dosing regimens should therefore be designed with consideration of the region-specific the probability of AQ over-dosing to reduce the risk of overdosing in the most at-risk populations.

## Abbreviations

ACT: artemisinin-based combination therapy AQ amodiaquine
AS-AQ: artesunate-amodiaquine CQ chloroquine
CV: coefficient of variation
DEAQ: desethylamodiaquine
DHA: dihydroartemisinin
FDC: fixed-dose combination
LoD: limit of microscopic detection
MDA: mass drug administration
MQ: mefloquine
PCR: polymerase chain reaction
PCT: parasite clearance times
PD: pharmacodynamic
PK: pharmacokinetic
PoC: periods of chemoprophylaxis
SP: sulphadoxine-pyrimethamine
WHO: World Health Organization
WWARN: Worldwide Antimalarial Resistance Network

## Ethics approval and consent to participate

Not applicable

## Consent for publication

Not applicable

## Authors’ Contributions

EMH & KK conceived the study. KBH calibrated and validated the model, and generated the results. All authors interpreted the results. KBH wrote the first draft of the manuscript. All authors gave critical input and contributed to the writing of the manuscript. All authors have read and approved the final manuscript.

